# Integrin α5β1 nano-presentation regulates collective keratinocyte migration independent of substrate rigidity

**DOI:** 10.1101/2021.03.08.434437

**Authors:** Jacopo Di Russo, Jennifer L. Young, Julian W. R. Wegner, Timmy Steins, Horst Kessler, Joachim P. Spatz

## Abstract

Nanometer-scale properties of the extracellular matrix influence many biological processes, including cell motility. While much information is available for single cell migration, to date, no knowledge exists on how the nanoscale presentation of extracellular matrix receptors influences collective cell migration. In wound healing, basal keratinocytes collectively migrate on a fibronectin-rich provisional basement membrane to re-epithelialize the injured skin. Among other receptors, the fibronectin receptor integrin α5β1 plays a pivotal role in this process. Using a highly specific integrin α5β1 peptidomimetic combined with nanopatterned hydrogels, we show that keratinocyte sheets regulate their migration ability at an optimal integrin α5β1 nanospacing. This efficiency relies on the effective propagation of stresses within the cell monolayer independent of substrate stiffness. For the first time, this work highlights the importance of extracellular matrix receptor nanoscale organization required for efficient tissue regeneration.

## Introduction

Collective cell migration is a fundamental biological process characterized by the coordinated movement of interconnected cells to achieve specific functions^1–3^. Basal keratinocytes collectively migrate to re-epithelialize a wound in the skin as the first step to re-establish tissue integrity^4,5^. At the onset of migration, keratinocytes deposit a provisional basement membrane that provides the necessary support for adhesion required for cell locomotion^6^. Among other components of this provisional matrix, fibronectin plays a pivotal role in supporting keratinocyte migration via the adhesion of α5β1 and αvβ1 integrins that are upregulated in re-epithelialization^7–9^. In contrast to αvβ1, which plays only a minor role in migration, α5β1 is required for efficient re-epithelialization, partially through the regulation of cellular traction forces^10,11^. In epithelial collective migration, each cell needs to coordinate cell-matrix traction forces with cell-cell stresses in order to propagate directed migratory cues to surrounding neighbours^2,12–14^. Previous work from our group and others have elucidated the many aspects of intercellular stress coordination governing collective cell migration ^12,13,15–17^, including the overall stress heterogeneity present in the migratory cell sheets, as well as the single molecular players involved in the mechanotransduction process^13,16^. Nevertheless, the role of extracellular matrix (ECM) adhesive sites in regulating such migratory coordination remains largely unknown.

Over the past two decades, it has become clear that nanometer-scale properties of the ECM strongly influence cell motility, in particular at the single cell level^18,19^. Previous work from our lab has highlighted the critical role of integrin lateral nanospacing in controlling single cell spreading, migration speed, and persistence^20,21^. Additionally, lateral adhesion spacing has been shown to regulate intracellular force generation in a manner consistent with the molecular clutch model^19^. In order to understand the role of ECM nanoscale organization in collective cell migration, we took a bottom-up approach by synthesizing nanopatterned hydrogels of discrete spacings, such that adhesive site organization could be precisely controlled. In order to investigate integrin α5β1 regulation of keratinocyte re-epithelialization, we utilized an engineered α5β1 integrin-specific peptidomimetic as the adhesive ligand (Fig. S1A). This specific peptide has a high binding affinity for α5β1 integrin (IC_50_ = 1.5nM), but orders of magnitude lower affinity for the other RGD-recognizing integrins, including αv-containing isoforms^22^. The advantage of using synthetic hydrogels lies in their ability to better mimic the physical properties of the ECM in the wound, as well as in their protein-repellent nature. With this approach, we could systematically address the role of cell-ECM interactions in cell monolayers by controlling both ECM stiffness and ligand density with nanometer precision. We were able to show that keratinocytes require an optimal integrin α5β1 spacing in order to efficiently coordinate their collective movement independent from ECM stiffness.

## Results

### Integrin α5β1 surface density controls keratinocyte migration efficiency

To investigate the role of integrin α5β1 in regulating keratinocyte collective migration, we fabricated gold nanopatterns on glass surfaces via block-copolymer micelle nanolithography (BCMN). BCMN allows for the precise modulation of ligand density at the nanometer level. The nanopatterns were covalently transferred onto polyacrylamide (PAA) hydrogels of ~ 23 kPa stiffness (approximately the stiffness of freshly wounded skin)^23,24^. The use of PAA hydrogels as culture substrate allows us to provide relevant substrate rigidity to cells and systematically control surface adhesion biochemistry due to the protein-repellent nature of PAA. Therefore, cells will only interact at nanoparticle sites, where highly specific integrin α5β1 peptidomimetics are linked, thereby allowing cell-surface adhesion via integrin α5β1 in a precisely controlled manner^25,26^ (Fig. 1A). In continuity with previous studies indicating cell-ligand length scale relevance for focal adhesion formation and cell motility, we modulated ligand density using three discrete inter-ligand distances: 35, 50, and 70 nm^20,21^.

**Figure 1:**
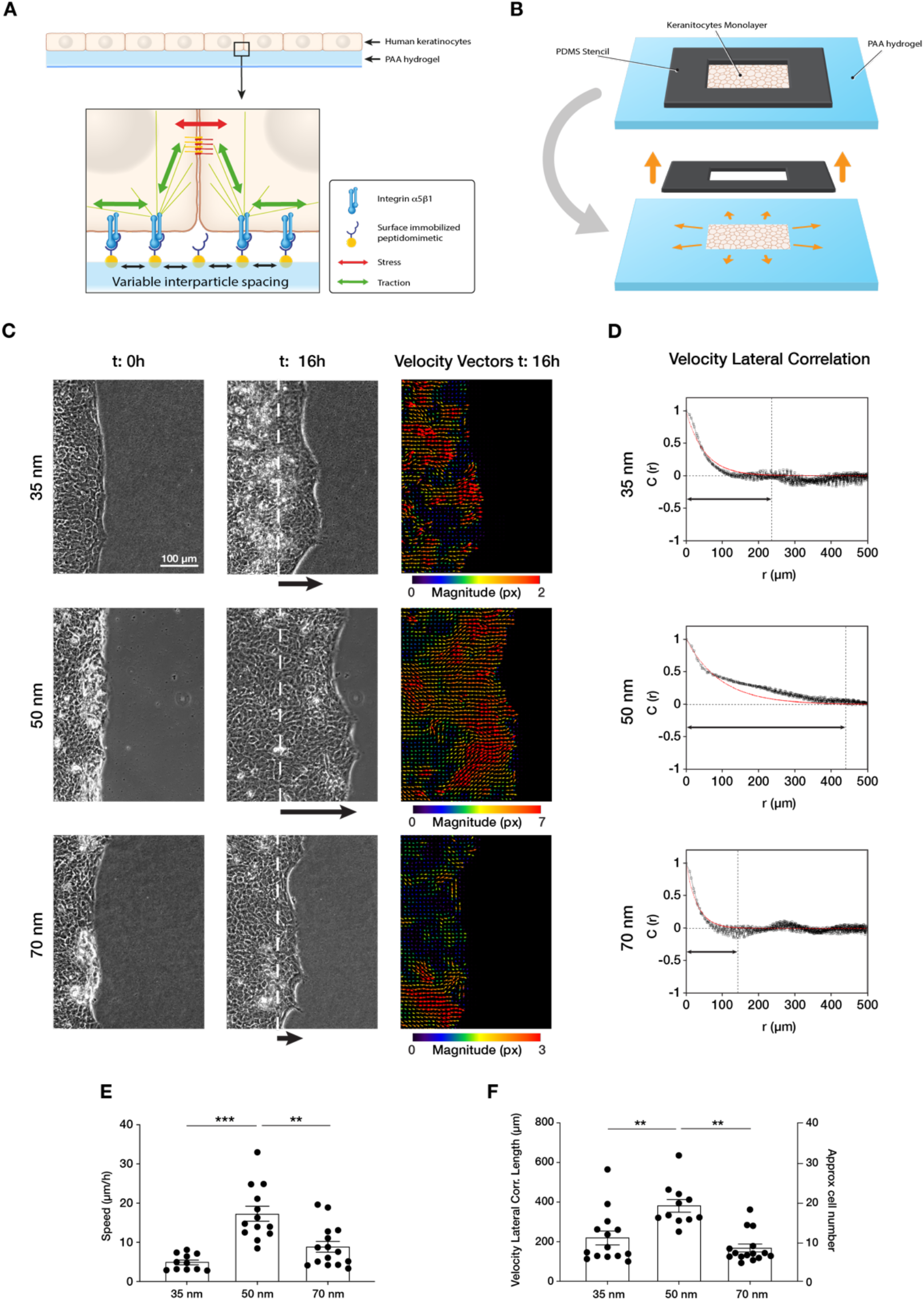
Efficient collective cell migration depends on integrin α5β1 ligand nanospacing. **A)** Keratinocyte monolayers coordinate their intra-/intercellular tractions/stresses on polyacrylamide (PAA) hydrogels nanopatterned with integrin α5β1 peptidomimetic. **B**) Schematic representation of the migration experiment setup. **C)** Representative images of time lapse experiments at t = 0 h (start) and t = 16 h (end), with corresponding velocity vectors plots in pixel (px) of keratinocyte sheet migration on hydrogels with 35, 50 and 70 nm integrin α5β1 ligand lateral spacing. The dotted white lines illustrate the initial starting point of the monolayers immediately following stencil removal, and the arrows indicate the direction of movement. **D)** Representative velocity lateral correlation length, C(r) where r = distance, curves of keratinocyte velocity vectors at t = 16 h. The quantification of migration speed **(E)** and the lateral correlation length **(F)** show an optimum at 50 nm integrin α5β1 ligand lateral spacing. Scatter plots show values with mean ± s.e.m. from at least three independent experiments. ** p < 0.01, *** p < 0.001 using a Mann-Whitney test.

Confluent monolayers of human immortalized keratinocytes (HaCaT) were cultured within a lateral confinement using a polydimethylsiloxane (PDMS) stencil placed onto the nanopatterned PAA hydrogels (Fig. 1B). Cell monolayer density was comparable among all three inter-ligand distances (Fig. S2). After PDMS stencil removal, cell sheet migration was triggered onto the empty regions of the gel, mimicking the collective migration process of basal keratinocytes post-skin wounding^4,5^. Quantifying cell sheet migration speed over culture time revealed a differential migration efficiency on the three conditions, with the highest speed observed at 50 nm (~ 20 μm/h) vs. slower speeds at 35 and 70 nm (~10 μm/h) (Fig. 1C, E, sup. movie 1, 2 and 3).

Next, we examined the extent to which ECM nano-presentation affects the directional migration of individual cells within the monolayer. Each cell in the monolayer coordinates its movement with its neighbours, and the length of such coordination can be quantified by comparing the lateral components (orthogonal to the direction of sheet migration) of the individual cell velocity vectors in the monolayer^13^. We thus used particle image velocimetry (PIV) to measure the velocity fields of migrating keratinocytes after 16 hours and calculated the lateral correlation length, C(r) (Fig. 1C). This revealed that keratinocytes migrating on 50 nm inter-ligand spacing most efficiently coordinate their movements, with an average correlation which extended over ~ 20 cells (~ 400 μm) (Fig. 1D, F). In contrast, keratinocytes on both 35 nm and 70 nm inter-ligand spacing were less able to coordinate their movements, only propagating their movements across a few cells (~ 200 μm) (Fig. 1D, F).

The density of integrin-ligand interactions alters cell spreading and migratory persistence through the regulation of focal adhesion maturation and dynamics in fibroblasts^18,20^. Since the correlation length of epithelial cell migration can be regarded as single cell migratory persistence, we analysed how integrin α5β1 lateral spacing influences focal adhesion dynamics (Fig. 2). After transient transfection of HaCaT cells with a mCherry-α-paxillin construct, we tracked focal adhesions and quantified their lifetime (sup. movies 4, 5, 6). Cells exhibited the highest ability to remodel their focal adhesions when migrating on 50 nm-spaced α5β1 ligands, resulting in the fastest turnover, or lowest lifetime. In contrast, on 35 and 70 nm inter-ligand spacing, focal adhesions exhibited longer lifetimes and therefore slower remodelling (Fig. 2A, B).

**Figure 2:**
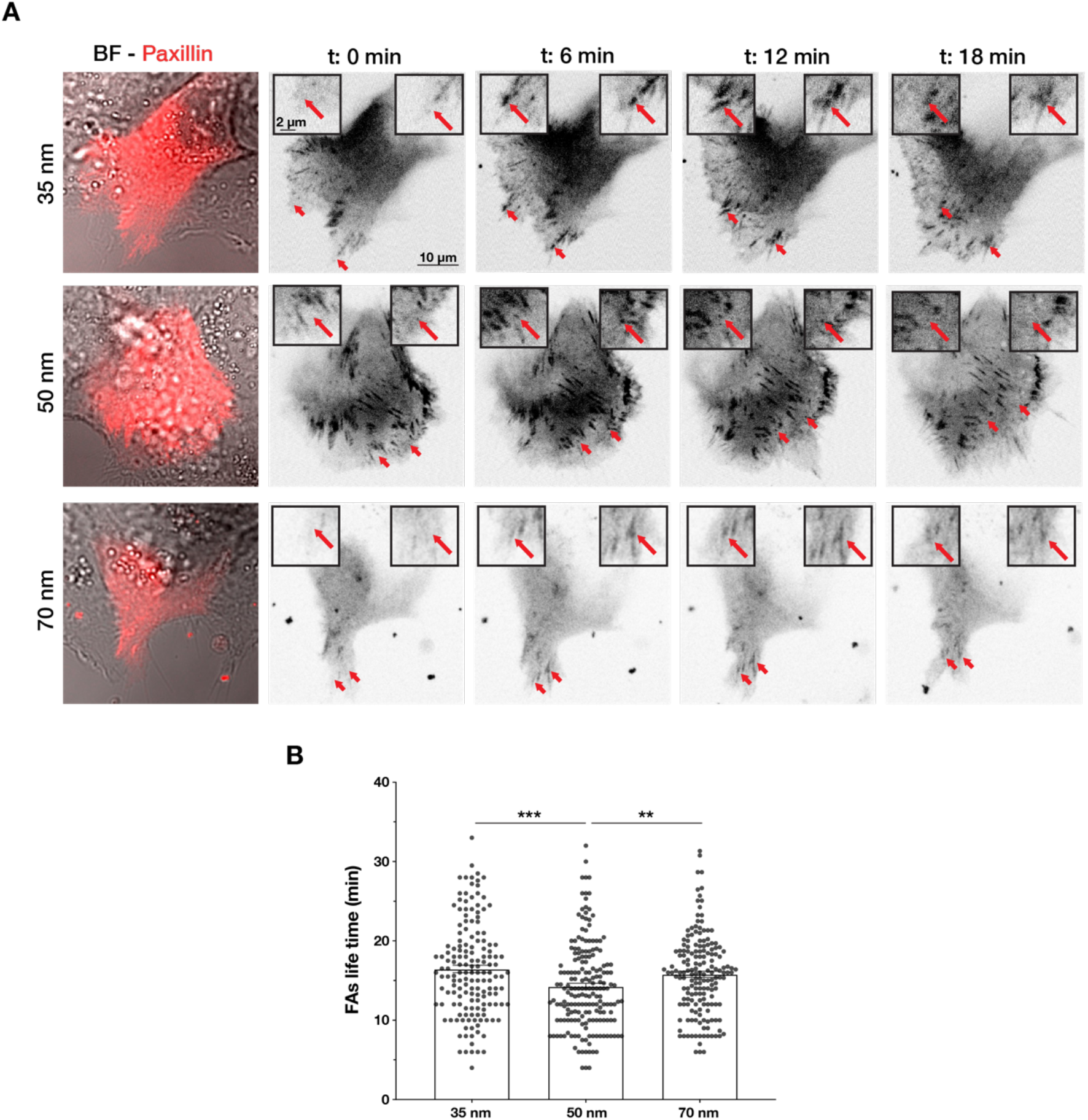
Integrin α5β1 lateral spacing governs focal adhesion dynamics. **A)** Representative still frames from time sequence over 18 minutes from transiently transfected keratinocytes with mCherry-α-paxillin migrating on 35, 50 or 70 nm integrin α5β1 ligand lateral spacing hydrogels. The red arrows and insets highlight the development of the same focal adhesion over culture time. Left column images are of BF = bright field overlaid with mCherry-α-paxillin (red) at t = 0. Other images are of mCherry-α-paxillin in B&W over culture time (t = 0, 6, 12, 18 min). **B)** Quantification of focal adhesion (FA) life time on the three different spacing surfaces. Column bars show mean values ± s.e.m of > 10 cells/condition, from at least three independent experiments. ** p < 0.01; *** p < 0.001 using a Mann-Whitney test.

Yet, α5β1 is not the sole ligand involved in keratinocyte migration. During wound healing, basal keratinocytes also interact with their provisional basement membrane using αv-containing integrin isoforms. We therefore examined whether migration was regulated in the same manner when engaging other integrins by using the c(RGDfK) peptide for cell attachment (Fig. S3A). c(RGDfK) allows for cell adhesion through a broader range of integrins, including αvβ1 and αvβ6, which are expressed by basal keratinocytes in the wound^8,22,27,28^. While c(RGDfK) still allows for α5β1 interaction, the affinity for this integrin subtype is lower compared to αv-containing isoforms (IC_50_ = 200nM)^22^(Fig. S1B). Interestingly, on c(RGDfK), keratinocytes exhibit faster migration on 35 nm inter-ligand spacing (vs. 50 nm on α5β1-presenting substrates), and correlation length does not change with receptor densities, which we showed to be biphasic for α5β1 (Fig. S3B, C). All together, these data reveal the existence of a specific optimal integrin α5β1 density that promotes faster focal adhesion dynamics and more efficient keratinocyte coordination, thereby leading to the most effective re-epithelialization process.

### Integrin α5β1 nanospacing controls force propagation in keratinocyte monolayers

Epithelial cells coordinate their movements as intercellular stress polarization increases, leading to collective cell migration through mechanotransductive processes^12,13,29^. Integrin nano-presentation in single cells has been shown to regulate not only the persistence of migration but also traction force generation^19,20^. Therefore, we sought to understand how integrin α5β1 density controls the amount of surface traction forces, and consequently, the intercellular stresses in the monolayer. To do this, we incorporated fluorescent beads into nanopatterned PAA hydrogels to perform traction force microscopy (TFM) on migrating monolayers. Surprisingly, this revealed a significant increase in monolayer tractions with decreasing integrin α5β1 ligand density (Fig. 3C, D), which did not follow the optimum migratory behaviour (velocity, correlation length) that we observed on 50 nm inter-ligand spacing (Fig. 1). We previously showed that the distance of stress propagation within the monolayer is crucial for collective cell migration, and that this is related to the levels of actomyosin contraction in individual cells^13,17^. Therefore, we were curious if integrin α5β1 lateral spacing differentially regulates the length scale of stress propagation in keratinocyte sheets, independent from the amount of traction forces generated.

**Figure 3:**
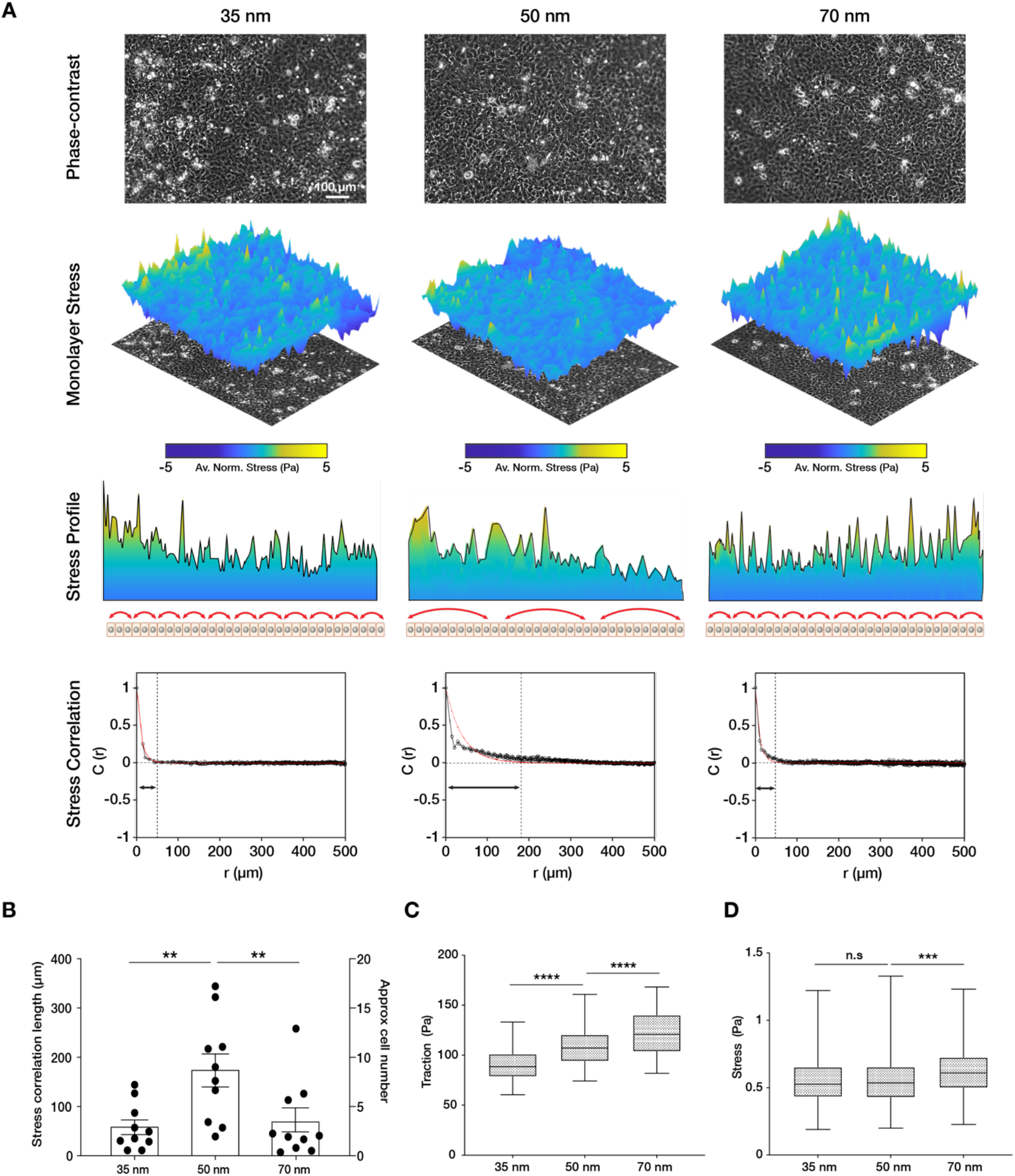
Traction force generation depends on integrin α5β1 ligand nanospacing. **A)** Representative phasecontrast images, monolayer stress plots and profiles, and spatial correlation curves from keratinocyte monolayers at the different integrin α5β1 ligand lateral spacings (35, 50, 70 nm). Stress correlation can be identified by inter-peak distance of the stress profile as schematically represented by the red arrows. Quantification of stress correlation length **(B)**, traction forces **(C)** and stresses **(D)** in the three spacing conditions. Scatter and box and whiskers plots show values and mean ± s.e.m from at least 4 independent experiments. n.s. = not significant; ** p < 0.01; *** p < 0.001; **** p < 0.0001 using a Mann-Whitney test.

To investigate this, we calculated the stress vectors using a force balance algorithm (monolayer stress microscopy - MSM)^12,17^ from the measured traction forces. We did this within confluent monolayers themselves in order to avoid any bias resulting from migration of the cell sheets (Fig. 3A). While the comparison of the absolute values of stress was in line with the traction forces, the spatial correlation length of the stress vectors was the largest on 50 nm inter-ligand distance (C(r) ~ 200μm) versus on 35 nm or 70 nm (C(r) ~ 50 μm) (Fig. 3A, B). To understand if this stress correlation length relationship was specific to α5β1 integrin, we performed MSM on c(RGDfK)-functionalized substrates. The results show no difference between the force correlation length on the three interligand densities, underlining the specific role of α5β1 in controlling the length of force propagation in keratinocyte monolayers (Fig. S3D).

In conclusion, the monolayer force correlation length was in line with the velocity correlation length in migrating keratinocyte sheets, suggesting that keratinocytes have an optimum integrin α5β1 density that best dictates the direction of intercellular stresses, and thereby the extent of force propagation in the monolayer.

### Integrin α5β1 lateral spacing overrides substrate stiffness and controls E-cadherin-mediated collective cell migration efficiency

It has been shown both *in vivo* and *in vitro* that collective cell migration is affected by substrate rigidity, with stiffer substrates generally enhancing migration efficiency^30–32^. Furthermore, it has been observed that focal adhesion dynamics increase with increasing substrate stiffness^33^. Therefore, we sought to understand the influence of substrate stiffness on the integrin α5β1 lateral spacing effect previously described on 23 kPa hydrogels. Migration and MSM experiments were performed within an *in vivo*-relevant range of tissue stiffness, i.e. on PAA hydrogels of 11, 23, 55, and 90 kPa,^24^ at the three inter-ligand spacings (35, 50, and 70 nm). Unlike on protein-coated surfaces^17^, keratinocytes were not able to form cohesive monolayers on the softer (< 11 kPa) or stiffer (> 90 kPa) nanopatterned surfaces, regardless of spacing. When comparing stiffness alone, higher substrate rigidity induced higher migration speed, as previously reported by others^30,31^ (Fig. 4A). Interestingly, we observed for each stiffness that cell sheet migratory speed was always more efficient when integrin α5β1 ligand lateral spacing was 50 nm vs. 35 or 70 nm (11 kPa: ~ 5 μm/h at 50 nm vs. < ~ 3 μm/h at 35 and 70 nm; 23kPa: ~ 17 μm/h vs. < ~ 9 μm/h; 55 kPa: ~ 15 μm/h vs. < ~ 10 μm/h; 90 kPa: ~ 14 μm/h vs. < ~ 9 μm/h) (Fig. 4A). The correlation of velocity vectors also confirmed a greater ability of keratinocytes to coordinate migration on the 50 nm inter-ligand spaced substrates (11 kPa: ~ 300 μm vs. ~200 μm; 23kPa: ~400 μm vs < ~ 200 μm; 55 kPa: ~ 400 μm vs. ~200 μm; 90 kPa: ~ 350 μm vs. < ~ 190 μm) (Fig. 4B). The extent of force propagation within the monolayers also confirms the higher force-propagation capability when integrin α5β1 ligand has a lateral spacing of 50 nm (Fig. 4C). MSM could not be carried out reliably on the stiff 90 kPa hydrogels due to limited bead movement caused by the specific surface functionalization, resulting in a high signal to noise ratio of the bead displacements. Nevertheless, speed and velocity correlation length observations on 90 kPa hydrogels show the 50 nm trend that we observe on the softer substrates.

**Figure 4:**
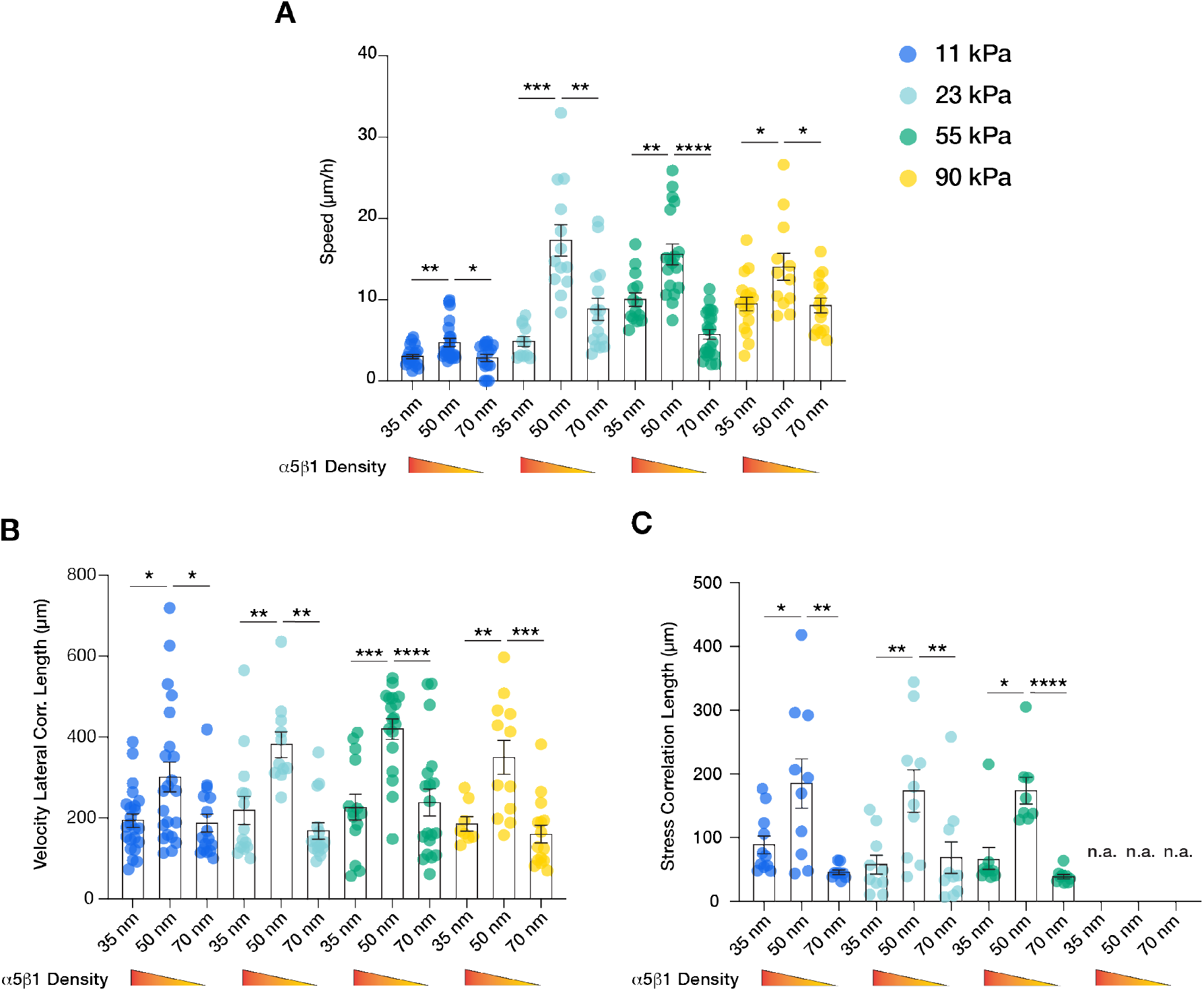
α5β1 ligand spacing outweighs substrate stiffness in collective cell migration. Quantification of the migratory speed **(A)**, velocity lateral correlation length **(B)** and stress correlation length **(C)** of keratinocyte sheets on different substrate rigidities (11 kPa - blue, 23 kPa - teal, 55 kPa - green, 90 kPa - yellow) and integrin α5β1 ligand lateral spacing (35, 50, 70 nm). Scatter plots show values and mean ± s.e.m from at least three independent experiments. n.a. = not applicable; * p < 0.05; ** p < 0.01; *** p < 0.001; **** p < 0.0001 using a Mann-Whitney test.

Finally, we sought to understand how integrin α5β1 ligand spacing directly regulates intercellular force transmission efficiency in keratinocytes. In epithelial sheets, the actin cytoskeleton physically connects cell-ECM adhesion structures with cell-cell adhesions (Fig. 1A). Between the latter, adherens junctions are responsible for maintaining the intercellular force continuum via E-cadherin-based homophilic interactions^16,30,34^. This mechanical continuum is crucial for coordinating collective migration by regulating intercellular pulling forces necessary for mechanotransduction pathways upstream of cell polarization^13^. We hypothesised that outside of the optimal 50 nm inter-ligand spacing of α5β1 peptide we observed, single keratinocytes would move counterproductively, thereby inhibiting monolayer motion. Thus, we inhibited E-cadherin interactions with the addition of a specific blocking antibody (DECMA-1) as previously shown^13^. When E-cadherin was inhibited, keratinocyte migration was enhanced on both 35 and 70 nm interligand spacing vs. 50 nm, as indicated by the distance of migration over culture time (Fig. 5A, B).

**Figure 5:**
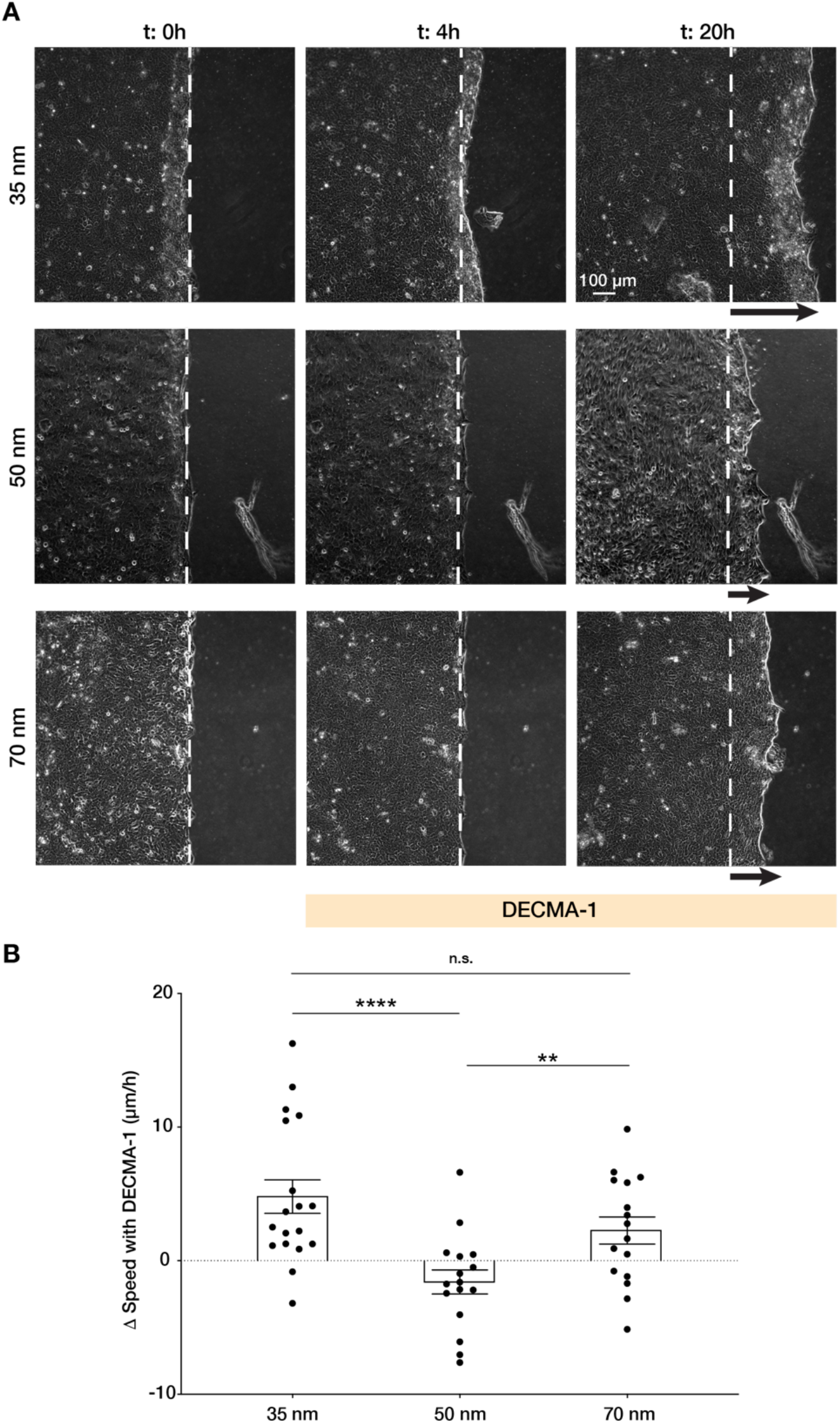
Integrin α5β1 spacing regulates intercellular force transmission via E-cadherin. **A)** Representative frames of keratinocyte sheet migration before (t = 0 - 4 h) and after (t = 4 - 20 h) the addition of E-cadherin blocking antibody (DECMA-1) in the medium. The arrows illustrate an approximation of the extent of sheet elongation. **B)** Quantification of the change in migratory speed in the presence of DECMA-1 vs. control. Scatter plots show values and mean ± s.e.m from at least three independent experiments. n.s. = not significant; ** p < 0.01; **** p < 0.0001 using a Mann-Whitney test.

Furthermore, when quantifying the change in sheet speed, Δv, after inhibition of E-cadherin vs. untreated cells, we observed migration speed was reduced on 50 nm vs. enhanced on 35 and 70 nm, indicating that the previously observed migration efficiency on 50 nm requires precise coordination between cell-cell and cell-ECM structures.

All together, these data indicate a vital role of integrin α5β1 lateral spacing in regulating keratinocyte collective migration. The optimal 50 nm lateral spacing outweighs the effect of substrate rigidity, as well as and regulates E-cadherin-mediated transmission of intercellular forces.

## Discussion

We show here that ligand nanospacing is integral to collective migration in keratinocytes, even outweighing the effects of substrate stiffness. Specifically, we found that 50 nm interligand distance is optimal for integrin α5β1, whereby collective cell migratory speed (Fig. 1C, E) and coordination were enhanced. At this distance, integrin α5β1 promotes the largest velocity correlation within the monolayer vs. at lower or higher ligand spacing (~ 400 μm on 50 nm vs. ~ 200 μm on 35 or 70 nm), indicating that cell movements are transmitted up to 20 cell lengths within the sheet. Consistent with enhanced migration, keratinocytes on this ligand spacing also exhibited the most dynamic focal adhesions, with short life times of ~ 14 minutes vs. ~ 16 minutes on 35 or 70 nm (Fig. 1D, F, Fig. 2). Quantification of the monolayer stress distribution revelated that this ideal inter-ligand spacing is not dependent on the absolute traction forces generated on the surface, but rather by the correlation length of stress vectors in the monolayer (Fig. 3A, B). This was also confirmed on hydrogels of varying Young’s moduli (11 - 90 kPa), with keratinocytes exhibiting enhanced migratory behaviour (faster and larger velocity and stress correlation lengths) at 50 nm spacing regardless of substrate stiffness (Fig. 4). This was particularly interesting because higher substrate rigidity is generally known to induce greater cellular traction forces, resulting in faster epithelial sheet migration^15,30,31^.

Each epithelial cell within a monolayer migrates in the direction of the maximal normal stress in a phenomenon called plithotaxis ^12,29^. In cell collectives, adherens junctions, which are connected to the actin cytoskeleton, are required for intercellular force transmission^16,30^. As the actin cytoskeleton is also connected to the ECM, this explains why we observed disrupted keratinocyte migration on 50 nm integrin α5β1 ligand lateral spacing when E-cadherin interactions were inhibited by addition of DECMA-1 antibody (Fig. 5). In contrast, at higher and lower ligand spacing, keratinocytes were able to migrate faster when we disturbed intercellular force transmission (Fig. 5). Therefore, we can conclude that outside of the optimal integrin α5β1 inter-receptor spacing of 50 nm, uncorrelated intercellular stresses slow keratinocyte sheet migration and result in inefficient collective movement (Fig. 6).

**Figure 6:**
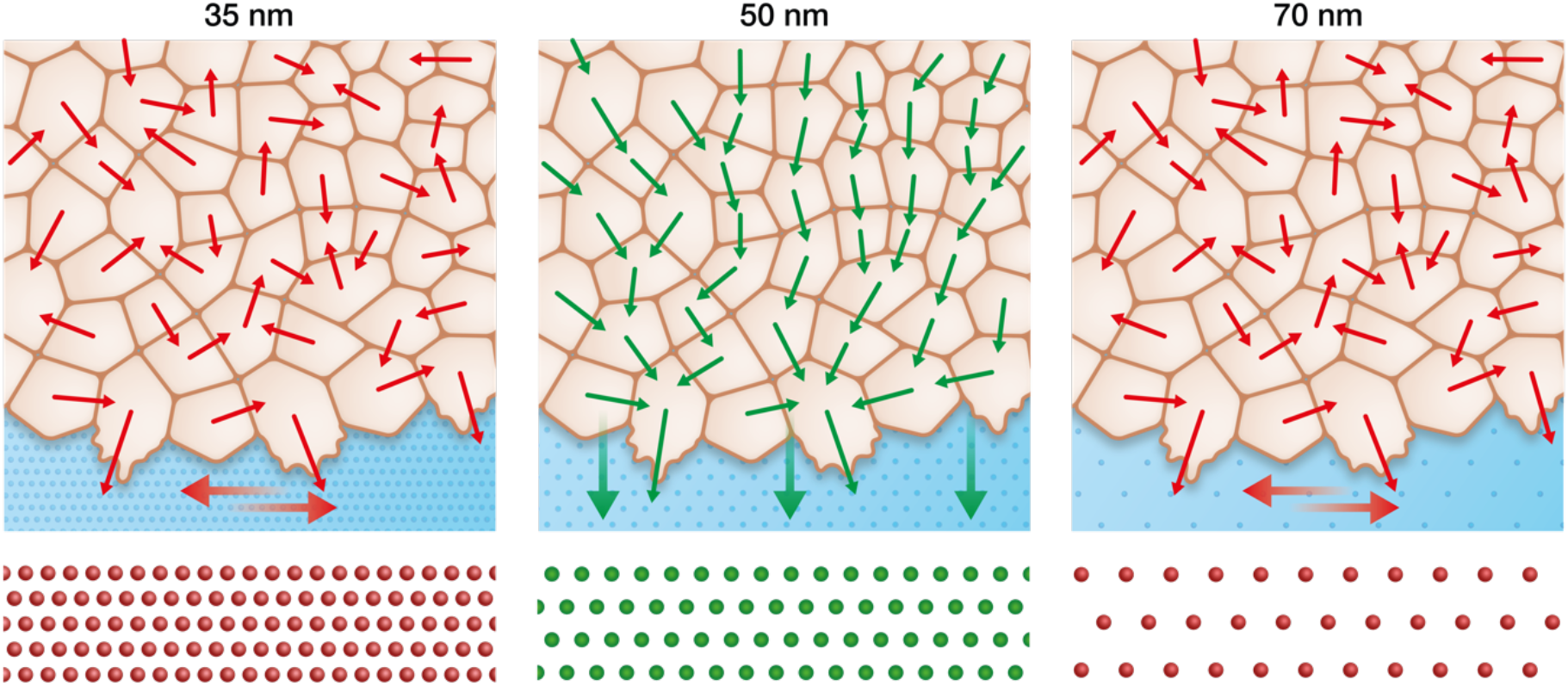
Keratinocytes require an optimum integrin α5β1 density to efficiently collectively migrate. In contrast to the optimal integrin α5β1 lateral spacing (green), lower and higher spacings (red) lead to uncorrelated single-cell movement and stress propagation. This results in inefficient collective behaviour, significantly slowing keratinocyte sheet migration.

It has previously been shown in single cells that integrin α5β1 regulates the generation of traction forces upon adhesion, whereas αv-class integrins mediate ECM rigidity sensing^11^. While we do observe faster collective migration with increasing substrate stiffness, as has been previously reported (Fig. 4A)^15,30,31^, α5β1 inter-receptor spacing outweighs stiffness effects alone. This could be attributed to integrin α5β1’s role in regulating traction force generation but not rigidity sensing. Indeed, higher substrate rigidity causes higher traction forces at the sheet edge at the outset of migration (removal of PDMS stencil), thereby transmitting higher net stress to the cell followers, ultimately resulting in more efficient collective migration^15^. This validates our data that 50 nm lateral spacing of integrin α5β1 is required for efficient force transmission independent from the net force exerted on the cells.

The severe phenotype acquired by murine animal models genetically lacking α5 or β1 integrin subunits has prevented our deep understanding into the roles specific integrin subtypes and organization play in wound healing^35,36^. With our bottom-up approach, we systematically identified the importance of integrin α5β1 density in coordinating force propagation within keratinocyte monolayers. How this inter-ligand optimal spacing can be understood in the *in vivo* context still requires further exploration, but our findings suggest that an integrin subtype-specific arrangement may be present in specific biological contexts that function through mechanosensitive pathways mediated by focal adhesions. Future studies will be required to address this phenomenon in order to more closely dissect the role of integrins and associated signalling pathways in the mechanobiology of wound closure.

## Contributions

J.D.R. and J.P.S. conceived the project. J.D.R. designed and performed all the experiments. J.L.Y. designed and prepared the nanopatterned hydrogels. J.W.R.W. and T.S. supported the technical execution of all the experiments. J.D.R., J.L.Y. and J.P.S. analysed and interpreted the data. H.K. provided the peptidomimetics. J.D.R. and J.L.Y. wrote the paper, and all authors discussed and commented on it.

## Acknowledgment

We thank the general support of the Max Planck Society and the grant from the Interdisciplinary Centre for Clinical Research within the faculty of Medicine at the RWTH Aachen University. We thank Adam Breitscheidel for his support with the graphic design.

## Methods

### Au-nanopatterned glass surfaces preparation

To obtain polyacrylamide nanopatterned surfaces, the desired Au-nanoparticle pattern was first obtained on 18 mm diameter glass coverslips (Carl Roth, Germany) as previously described^20,23^. Briefly, the coverslips were cleaned in piranha solution (3 H_2_SO_4_:1 H_2_O_2_), rinsed thoroughly with MilliQ water and stored in a dust-free environment until further use. Au-micellar solution was obtained first by dissolving a diblock copolymer polystyrene-b-poly(2-vinylpyridine) (Polymer Source) in o–xylene and second by loading with HAuCl_4_ **·** 3H_2_O (Sigma Aldrich) according to the specific parameter L = n[HAuCl_4_]/n[P2VP]. The prepared micellar solution was used to spin coat the clean coverslips which were subsequently treated with argon-hydrogen plasma (90% Ar/10% H_2_) in a Tepla PS210 microwave plasma system (PVA Tepla, Germany) for 45 minutes at 200 W and 0.4 mbar in order to remove the polymer. Different ratios of polystyrene/poly(2-vinylpyridine) units (288/119 or 501/323) and spin speeds (3000 – 8000 rpm) were employed to obtain 35, 50 and 70 nm interparticle spacing. Nanopatterned glass surfaces were then evaluated by scanning electron microscopy (SEM) (Carl Zeiss, Germany) as previously described^20,23^. Interparticle spacing and overall order was quantified with the K-Nearest-Neighbours algorithm implemented by an in-house script written in ImageJ software. The k-nearest neighbours (k = 6 for ordered particles) was estimated for more than 600 particles per sample.

### Transfer of Au-nanopatterns onto polyacrylamide hydrogels

To be able to transfer the gold nanoparticles from the glass surface to the hydrogels while preserving their distribution, the nanopatterned coverslips were incubated with 0.5 mg/ml N, N’-bis(acryloyl)cystamine (Thermo Fisher) in ethanol for 1 hour at room temperature, washed with ethanol and dried with nitrogen stream. Afterwards, hydrogels were formed by pipetting a mixture of acrylamide (Bio-Rad), bis-acrylamide (Bio-Rad), 0.003% Tetramethylethylenediamine (TEMED, Bio-Rad), and 0.03 % Ammonium persulfate (Sigma) diluted in phosphate-buffered saline (PBS) between glutaraldehyde-activated glass bottom dishes (MatTek) and AuNP-functionalized coverslips. The following ratio of acrylamide/bis-acrylamide were employed to reach different hydrogel Young’s moduli: 10%/0.07% for 11 kPa; 7.5%/0.2% for 23 kPa; 12%/0.6% for 55 kPa; 12%/0.3% for 90 kPa^37^. Hydrogels were allowed to swell for at least 72 hours in PBS to facilitate the detachment of the coverslips and the transfer efficiency was evaluated by SEM of both the glass and hydrogels surface. The obtained hydrogels were sterilized with 30 minutes ultraviolet light irradiation before their employment as cell culture substrates.

### Cell culture

Human immortalized keratinocytes (HaCat) were cultured with Dulbecco’s Modified Eagle Medium (DMEM, Gibco, Thermo Fisher Scientific) supplemented with 10% Fetal Bovine Serum (FBS, Sigma Aldrich) and 1% Penicillin-Streptomycin (Gibco, Thermo Fisher Scientific) at 5% CO_2_ and 37 °C. To passage and perform experiments, cells were detached with 10 minutes incubation with 5 mM EDTA solution in PBS followed by the incubation with 0.05% Trypsin/EDTA (Gibco, Thermo Fisher Scientific).

### Migration experiments

To perform cell migration experiments, nanopatterned hydrogels were incubated for two hours with 25 μM customized thiolated c(RGDfK) peptide (PSL GmbH, Heidelberg, Germany)^38^ (Fig. S1B) or with thiolated integrin α5β1 peptidomimetic^39–41^ (Fig. S1A) to allow for cell adhesion. Bovine serum albumin-coated polydimethylsiloxane (PDMS) stencils were employed to horizontally confine the cells on the surface and obtain a confluent monolayer of approximately 3 x 6.5 mm. Cells were seeded with a density of 4500 cells per mm^2^, allowed to adhere and to reach confluency for approximately 12 hours at 5% CO_2_ and 37 °C, and triggered to migrate with the lift-off of the confinement. Cell migration experiments were carried out either inside a standalone incubator or within an incubator staged over an Axio Observer 7 (Carl Zeiss, Germany). Images were acquired every 10 minutes at multiple positions using an automated stage controlled by Zen software (Carl Zeiss, Germany). For E-Cadherin blocking experiments, cell monolayers were allowed to migrate for at least four hours before exchanging the medium with pre-warmed medium containing 10 μg/ml E-cadherin blocking antibody (clone DECMA-1, Millipore MABT26).

### Traction force and monolayer stress microscopy

Traction force and monolayer stress microscopy (TFM, MSM) were performed as previously described^13,17^. 1 μm fluorescent carboxylate-modified polystyrene beads (Sigma) were mixed in the polyacrylamide solution before polymerization and allowed to reach the hydrogel surface by gravity during gelation. Cell-induced bead displacement vectors were calculated by comparing the picture with relaxed beads (cell-free gel after trypsinization) from the picture acquired with cells using the particle image velocimetry (PIV) plugin of ImageJ. Traction forces were calculated from these vectors using the Fourier transform traction cytometry plugin (FTCC). Average normal stress vectors within the monolayer were calculated using the traction force information using a force balance algorithm in MATLAB (MathWorks) as formulated elsewhere^17^. The force propagation within the monolayer is characterized by the force correlation length, which is the length-scale of the following spatial autocorrelation function:

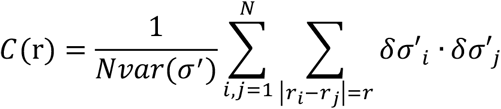

where *δσ*’ are the local deviation of the average normal stress at position from its spatial mean *σ*’ and var(σ’) is its variance. The correlation length of the stresses was determined at the point where the function was equal to 0.01^17^.

### Focal adhesion lifetime quantification

To be able to visualize focal adhesion dynamics and quantify their lifetime, HaCaTs were transfected with mCherry-α-Paxillin-C1 construct kindly provided by Dr. Eli Zamir^42^. Transient transfection was performed following the Fast-Forward protocol provided by the Attractene transfection reagent kit (Qiagen, Germany). Briefly, 0.1 μg of the plasmid per sample was allowed to react with the Attractene reagent for 15 minutes at room temperature and then mixed with the cell suspension for cell seeding within the PDMS stencil placed on the hydrogel. After 12 hours, cell migration was initiated and mCherry signal was acquired every two minutes using an LSM 880 confocal microscope (Carl Zeiss, Germany) equipped with a 40X longdistance water immersion objective with a numerical aperture of 1.1. The obtained time-lapse videos were analysed using Imaris image analyses software (Bitplane, Oxford Instrument) automatically tracking each focal adhesion in the cells and quantified its life time.

### Quantification of velocity vectors and correlation length

Velocity vectors and their correlation length, which quantify the ability of cells to coordinate their movements, was determined as previously described^13^. Briefly, time-lapse videos obtained using phasecontrast microscopy and two consecutive images with an interval of 10 minutes were used to calculate velocity vectors using the PIV plugin of ImageJ software. In PIV analyses, each image was broken down in 32 x 32 pixels windows for comparison and a two component *(i,j)* velocity vector was assigned to the center of the window, namely lateral *(U_ij_*, perpendicular to the direction of monolayer migration) and axial *(V_ij_*, along the monolayer migration). The fluctuations of the lateral component of the vectors (*v_ij_*) were determined as follows:

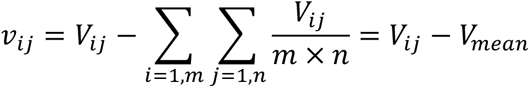

where *V_mean_* is the mean lateral velocity. The lateral velocity correlation function was formulated as following:

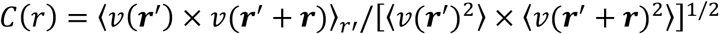

where *〈...〉* symbolizes the average, **r** is the vector of coordinate (*i,j*) and *r* is the norm of **r** defined as r = ||**r**||. Similar to the force correlation length, the lateral correlation length was determined at the point where the function was equal to 0.01^13^.

### Quantification of cell density

To quantify keratinocyte cell density on the different substrates, the monolayers were fixed for 10 minutes using 4% paraformaldehyde in PBS for the same conditions as in migration experiments. Upon permeabilization using 0.3% Triton X-100 for 2 minutes, the monolayers were stained for DNA using DAPI. The number of nuclei were counted using ImageJ software (NIH) in immunofluorescent imaged obtained at 10x magnification.

### Statistics

Statistical tests and graphics were performed using GraphPad Prism software, choosing parametric or not parametric tests after evaluating the normality distribution of the data. Sample conditions and the specific test used for each data set are indicated in corresponding figure captions.

