## Supplementary material for "Integrin α5β1 nano-presentation regulates collective keratinocyte migration independent of substrate rigidity": Fig. S1

### Supplementary Figures

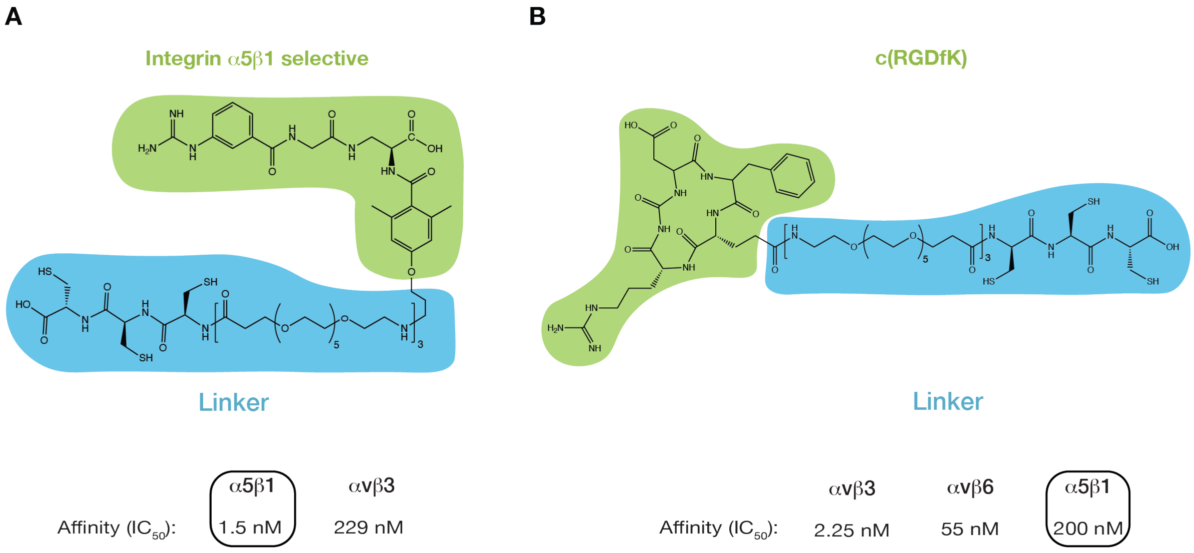


**Figure S1: Chemical structures of integrin α5β1 selective peptidomimetic (A) and c(RGDfK) (B).** Highlighted in green are the integrin-selective moieties of the peptides and in blue the functionalization required for interaction with the gold nanoparticles. At the bottom are the integrin subtype-specific affinity values as reported elsewhere[22,25].
