## Supplementary material for "Integrin α5β1 nano-presentation regulates collective keratinocyte migration independent of substrate rigidity": Fig. S2

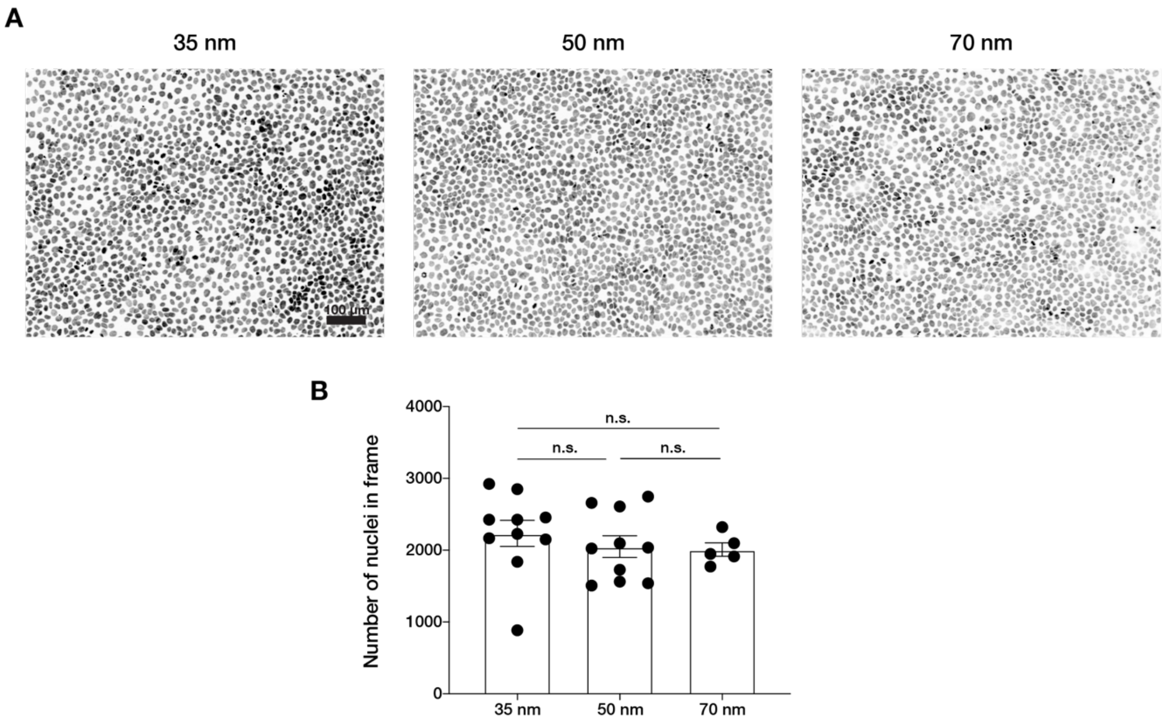


**Figure S2: Keratinocyte monolayer density is not affected by integrin α5β1 nanopatterning. A)** Representative immunofluorescent images of keratinocyte nuclei (stained with DAPI) in monolayers obtained on 35, 50 and 70 nm inter-ligand spacing. **B)** Quantification of number of nuclei per frame shows no significant differences in cell density between the different conditions. Column bars show mean ± s.e.m. from at least three independent experiments. n.s. = not significant using an unpaired t-test.
