## Supplementary material for "Integrin α5β1 nano-presentation regulates collective keratinocyte migration independent of substrate rigidity": Fig. S3

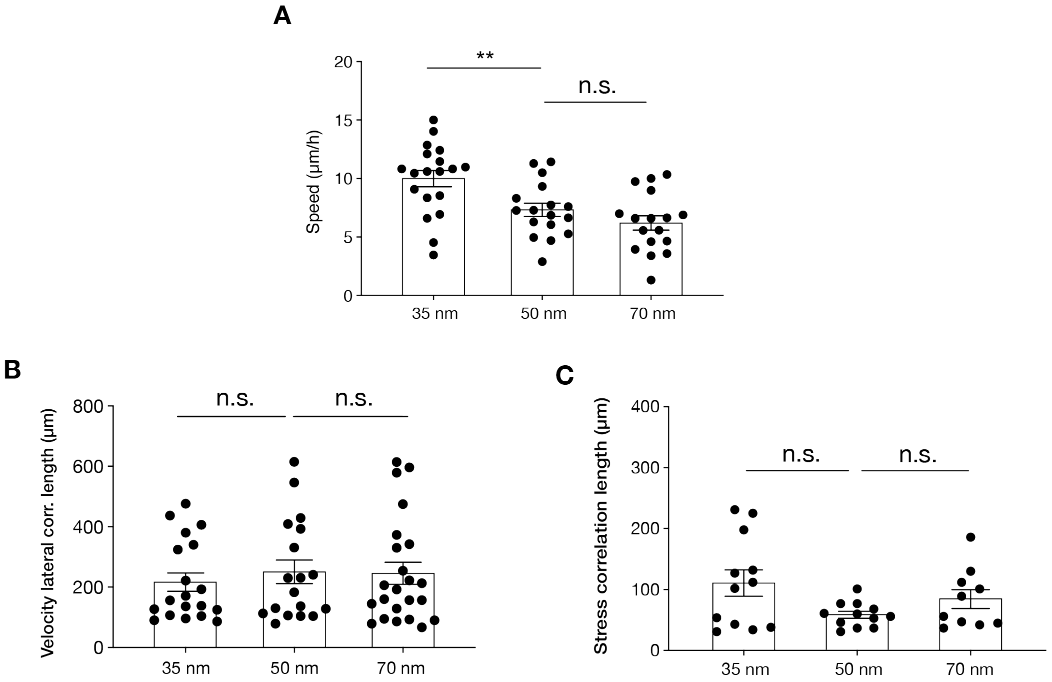


**Figure S1: Keratinocyte collective behaviour on nanopatterned c(RGDfK) surfaces. A)** Quantification of keratinocyte sheet migration speed, velocity lateral correlation length **(B)** and stress correlation length **(C)** on 35, 50 and 70 nm inter-receptor spacing. Scatter plots show individual values and mean ± s.e.m. from at least three independent experiments. ** $p < 0.01$, n.s = not significant using a Mann-Whitney test.
